# Transcription activation of early human development suggests DUX4 as an embryonic regulator

**DOI:** 10.1101/123208

**Authors:** Virpi Töhönen, Shintaro Katayama, Liselotte Vesterlund, Mona Sheikhi, Liselotte Antonsson, Giuditta Filippini-Cattaneo, Marisa Jaconi, Anna Johnsson, Sten Linnarsson, Outi Hovatta, Juha Kere

**Author notes:** These authors contributed equally (Virpi Töhönen, Shintaro Katayama).

## Abstract

In order to better understand human preimplantation development we applied massively parallel RNA sequencing on 337 single cells from oocytes up to 8-cell embryo blastomeres. Surprisingly, already before zygote pronuclear fusion we observed drastic changes in the transcriptome compared to the unfertilized egg: 1,804 gene transcripts and 32 repeat elements become more abundant, among these the double-homeobox gene *DUX4*. Several genes previously identified as *DUX4* targets, such as *CCNA1, KHDC1L* and *ZSCAN4*, as well as several members of the *RFPLs, TRIMS* and *PRAMEFs^1^* were accumulated in 4-cell stage blastomeres, suggesting *DUX4* as an early regulator. In the 8-cell stage, we observed two distinct cell types – a transcript-poor cell type, and a transcript-rich cell type with many Alu repeats and accumulated markers for pluripotency and stemness, telomere elongation and growth. In summary, this unprecedented detailed view of the first three days of human embryonic development reveals more complex changes in the transcriptome than what was previously known.

---

The study of early human development has been based on a small number of samples, often pooled, due to the rarity of material, and due to methodological reasons on selected genes for profiling with the subsequent risk of masking differences in gene expression between blastomeres^2-4^. In the present study, we sought to overcome these earlier limitations to obtain a more detailed view of human embryo development. To our knowledge this is the first single cell sequencing analysis investigating global gene expression within cells, individually isolated from cleavage stage embryos during the first 3 days of human development. A total of 337 oocytes, zygotes and blastomeres from 4- to 10-cell embryos were collected from clinics in Sweden and Switzerland (ethical permissions and legislation in Online Methods). To investigate global gene expression, we used single-cell tagged reverse transcription (STRT), a highly multiplexed method for single-cell mRNA-sequencing that allows molecule counting^5^. To test the validity of our findings, we performed sequencing on single cells using the Tang *et al* method^6^. All results were replicated in independent cells and sequencing experiments. The cells were collected into individual wells on a 96-well plate, reverse transcribed, barcoded, pooled and sequenced as one library (Online Methods, Supplementary Table 1 and Supplementary Data 1). In human, mitotic cell divisions occur in the absence of cellular growth that divides the volume of daughter blastomeres^7^ (Fig. 1a) and reduces the amount of transcripts per cell (Fig. 1b, supported by Dobson et al^8^). Therefore, in this study we used internal control mRNAs to estimate absolute mRNA count per cell to compensate for the reduction in transcript number.

**Figure 1.**
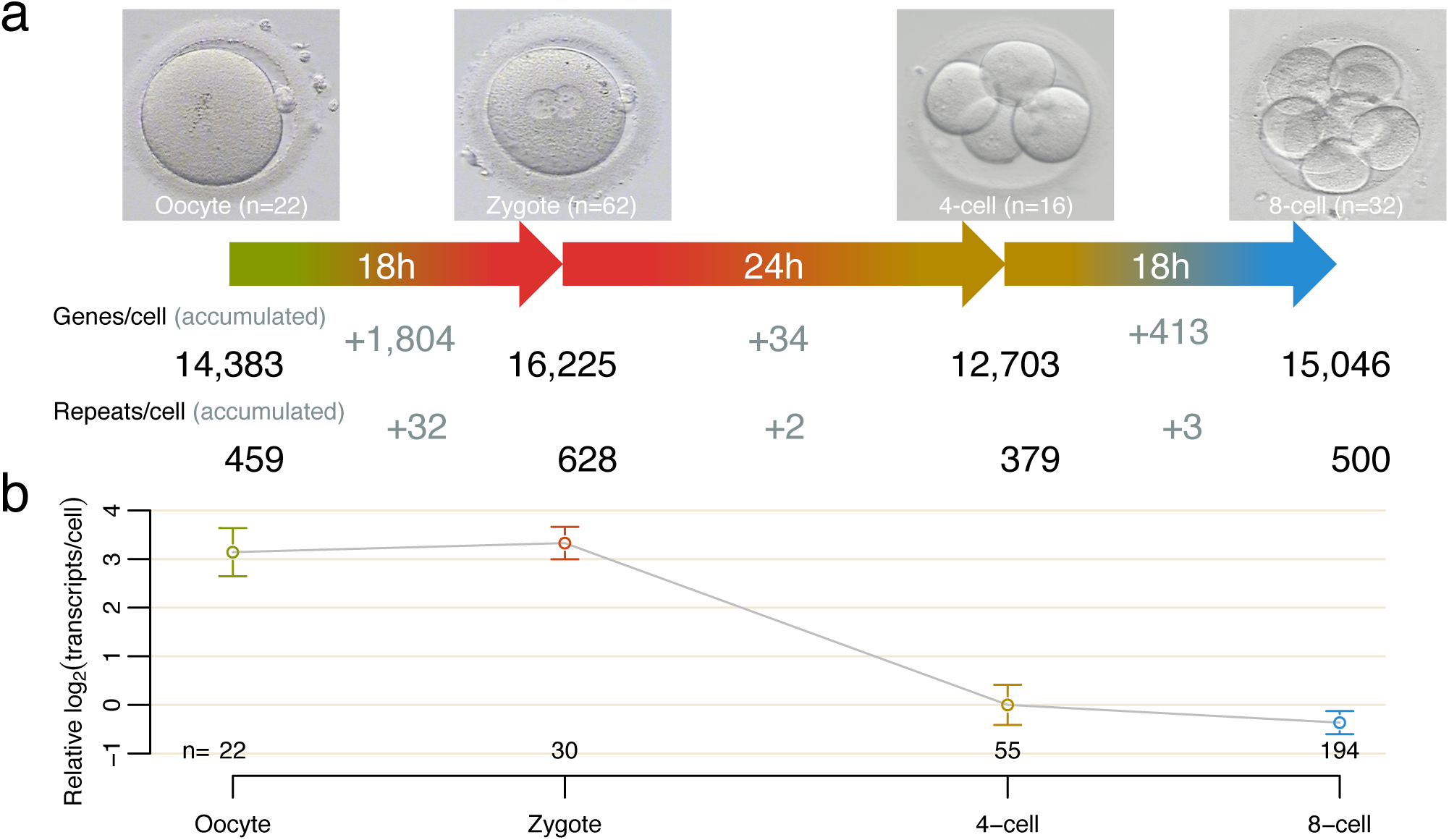
Transcriptome dynamics in human preimplantation development. a) Photomicrographs showing the morphological features of the four developmental stages studied. Starting from left is a metaphase II oocyte with an extruded first polar body; fertilization will give rise to the zygote, determined by confirmation of the male and female pronuclei containing haploid sets of chromosome; 4-cell stage embryo on day 2; and 8-cell embryo on day 3 of development. The indicated times between the stages are approximations for human embryo cleavages^35,36^. The numbers in the photos show the total number of oocytes, zygotes and embryos included in this study. The numbers below indicate the total gene and repeat number per cell, as well as the increase in transcripts between stages. Detected genes and repeats had more than eight reads in at least one sample. b) Detected transcriptome dynamics during the first 3 days of human preimplantation development. The numbers for each embryonic stage indicate individual oocytes, zygotes and blastomeres that were used for the estimation of relative transcript counts, using 4-cell stage blastomeres as reference. Embryos with three to five blastomeres were classified as 4-cell stage and six-to ten cell blastomeres as 8-cell stage. Vertical bars denote 95% confidence intervals. After fertilization an increase in available polyA transcripts was detected. In the subsequent two cleavages, we observed 8-fold reduction in transcript count between zygote and 4-cell stage, indicating both active (actual mRNA degradation) and passive (mRNA content division between daughter blastomeres) decrease.

We first compared transcript profiles in oocytes and zygotes, as they are stages just before and after fertilization. Previous studies using microarrays based on traditional assumption of equivalent transcriptome distributions have shown very little difference between oocyte and zygote stages^8^. We detected 1,804 genes and 32 repeat elements accumulated at the zygote stage (Fig. 2a, 2b and Supplementary Data 2) of which 44 genes were specifically detected only in zygotes (Fig. 2b and Supplementary Table 2). The most abundant among the zygote-specific genes was a transcript originating from the complex *D4Z4* region located on chromosomes 4q and 10q (Fig. 2c), which contains the *DUX2, DUX4* and *DUX4-like* genes. *DUX4* is a double homeobox-containing transcription factor expressed in a small number of cells in human muscle, testis, ovary and heart^9^.

**Figure 2.**
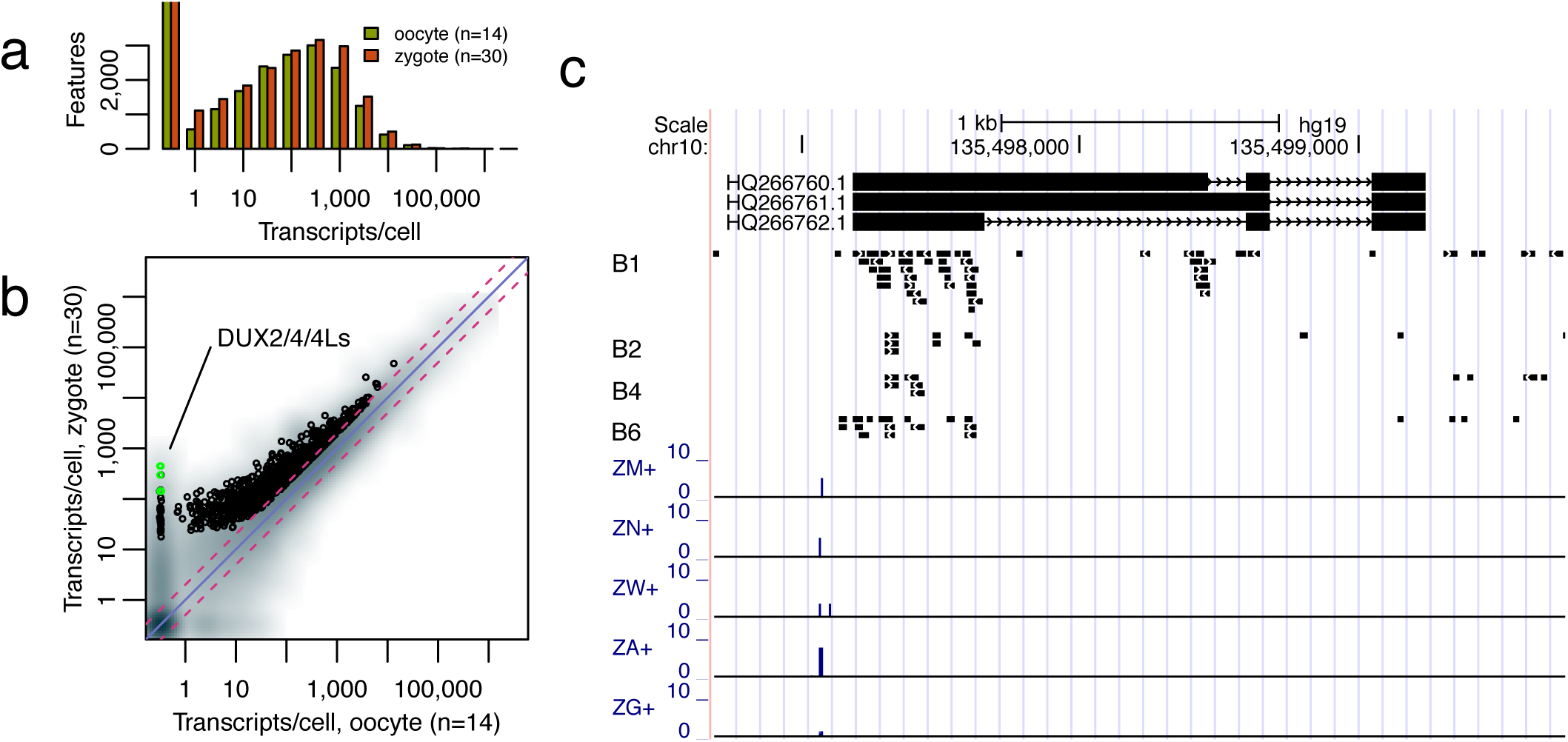
Transcriptome changes during the oocyte-to-zygote transition. a) In oocyte-to-zygote comparison on L233 library, the transcriptome increases in both the total transcript number and the numbers of different transcripts detected. b) Comparison of mRNA molecular count per cell per transcript on L233 library. Several of the observed accumulated transcripts in zygotes are known key developmental genes as well as genes not previously implicated in early human development. Particularly interesting are the small number of genes detected only in the zygote. Among these are transcripts of the *DUX4* gene from the highly repetitive D4Z4-region. No genes were detected as significantly downregulated in the transition from oocyte to zygote. Pink dashed lines = fold-change thresholds; 2-fold up- or down-regulation. Diagonal purple line = equal numbers of transcripts/cell between compared stages. c) Given in the figure is the organisation of the D4Z4 repeat element on chr10:135,496,683-135,499,746 at 10q26.3 (hg19 reference genome); +/-500bp of HQ266762, identified as DUX4-s by Snider *et* al^9^. Bottom five tracks show the strand-specific STRT signals observed in five representative zygotes on the forward strand. We used the Tang method for single cell sequencing^6^ to confirm the *DUX4* expression in zygotes (the method differs from STRT in that it is not strand specific or enriched for 5’UTR of transcripts). The four human zygotes sequenced are in the middle tracks, called B1, B2, B4 and B6. In addition, we confirmed the expression of both full length and short version of *DUX4* in a human zygote by Sanger sequencing (data not shown).

In our data, out of the 25 complex repeats that increased at zygote stage, 13 belong to *ERVs* and seven to *LINE* families^10^ (Supplementary Table 3). In recent years, retrotransposons such as ERV (Endogenous Retroviruses) and LINE (Long Interspersed Elements) have been shown important for early murine embryo development^10-12^. The observed change in the transcriptome profile in zygotes compared to oocytes may be due to active transcription caused by genome-wide demethylation occurring after fertilization^13^. Since polyA tail is a prerequisite for STRT sequencing, other known processes affecting the polyadenylation may cause the observed change. Such processes are cytoplasmic polyadenylation of maternally-derived transcripts, that has been shown as enriched in murine oocyte to zygote transition^14^, or recruitment of translationally inactive mRNAs stored in subcortical ribonucleoprotein particles in oocytes^15^.

We next addressed characteristics of the transition from zygote to 4-cell stage. The very first stages of embryo development are believed to be transcriptionally silent^16^ and thus translation and degradation of maternally inherited transcripts is required for successful maturation and completion of embryo genome activation (EGA)^17^. Multiple transcripts decreased from zygote to the 4-cell blastomeres (Fig. 3a and 3b) probably reflecting degradation of maternal RNA taking place after fertilization as well as passive reduction by cleavage divisions. The transcripts that increased during the same period consists of 34 genes and 2 repeat elements not previously attributed to human preimplantation development (Fig. 3b and Supplementary Data 2). In support of DUX4 as a developmental regulator, shown to both repress and activate specific gene transcription^1,18^, we found a total of 25 out of 34 genes recently identified as targets of DUX4 overexpression in human myoblasts^1^ increased between zygote and 4-cell stage (Supplementary Table 4). Furthermore, we find DUX4 activated repeat element *ERV-MaLR* accumulating during the zygote to 4-cell transition. Among the target genes detected at 4-cell stage we found *KHDC1L*, the cell cycle regulator *CCNA1* and *ZSCAN4* shown to be expressed in early embryos and involved in telomere maintenance and genomic stability^19,20^, as well as several members of the *RFPLs, PRAMEF* and *TRIM* families (Fig 3b and 3c). At least 8 genes from the *PRAMEFs* (Preferentially Expressed Antigen in Melanoma Family) were expressed (Supplementary Note). While the *PRAMEF* genes have been recognized in cancer, they have not previously been reported in early embryo development. A role for DNA methylation in *PRAMEF* gene regulation, with increased expression upon hypomethylation has been suggested^21^. Expression of *DUX4* (in individual zygotes) and of *ZSCAN4, CCNA1, KHDC1L, FRG2* and *DIO2* (in 4-cell human embryos) were further confirmed by PCR and Sanger sequencing (data not shown). Taken together, our data suggest *DUX4*, specifically expressed in the zygotes, as a novel embryonic transcription factor contributing to the first wave of transcription in 4-cell stage human embryo.

**Figure 3.**
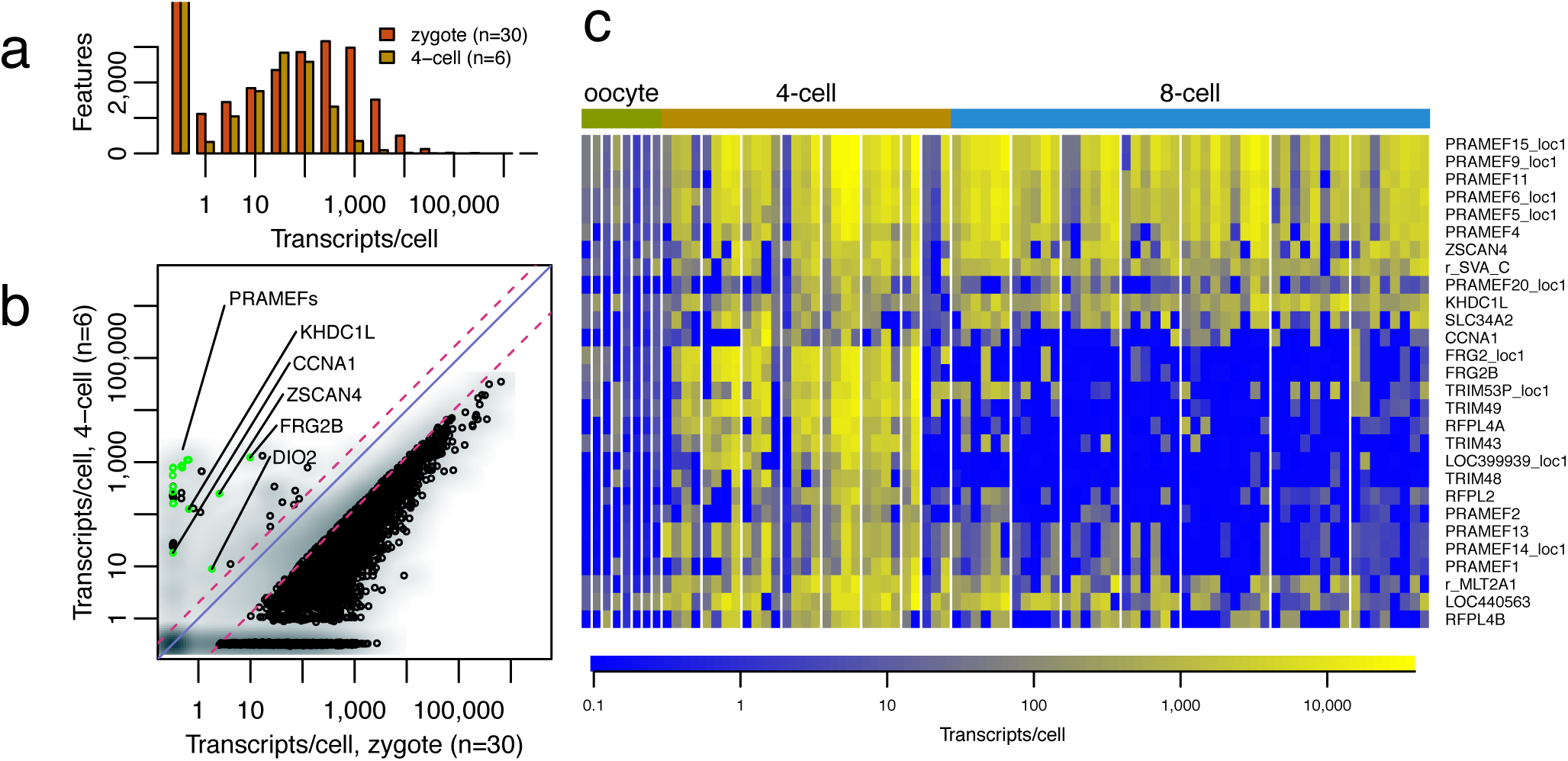
Transcripts differently regulated during zygote-to 4-cell transition. a) The comparison of mRNA molecular count per cell on L233 library showed reduced expression levels. This is observed as a shift of the overall peak of transcripts and a reduced area of expressed gene number in 4-cell blastomeres compared to zygotes. b) We confirmed expression of a subset of genes (highlighted) by PCR and Sanger sequencing of PCR products using separate 4-cell mRNA libraries. Pink dashed lines = fold-change thresholds, 2-fold upregulation or 8-fold downregulation. The diagonal purple line = equal numbers of transcripts/cell between the two compared stages. c) Heat map showing the genes accumulated at 4-cell stage on L185 library. Columns are cells separated by embryo. The individual cells within an embryo are sorted by transcript count/cell; with transcript-rich cells to the right. The majority of the accumulated genes are identical to DUX4 target genes^1^ (Supplementary Table 4). The bottom panel displays genes accumulated at 4-cell stage and then reduced. The top panel displays genes that stay expressed at the same level through 8-cell stage. Only one representative for each of the multi-copy genesis shown; Supplementary Data 2 includes all copies. The tripartite motif present in the TRIM (Tripartite motif containing protein) family proteins includes RING, B-box, and coiled-coil domains. Two additional transcripts detected, *RFPL4B* and *RFPL2*, also code for RING domain containing proteins implicated in a variety of biological processes such as the regulation of differentiation, development and oncogenesis (NCBI, conserved domains). Little or no expression of these genes was detected in the human oocytes, indicating that they were accumulated following fertilization.

The major embryonic genome activation is thought to occur between 4-and 8-cell stage of development^16^. According to PANTHER-based ontology analyses, the up- and down-regulated transcripts detected during this transition were involved mainly in transcriptional and translational activity, in cell cycle regulation, protein processing, transport and cell communication (Fig. 4a, 4b and Supplementary Data 2). When investigating global gene expression, we discovered statistically significant up-regulation of Y-chromosomal *ZNF100, RPS4Y1* and *DDX3Y*, indicating paternal genome activation already at the 4-8 cell stage in male embryos. In putative female embryos a corresponding statistically significant upregulation of *SOX4, KIF4A* and *DDX3X* was observed (Fig. 4c).

**Figure 4.**
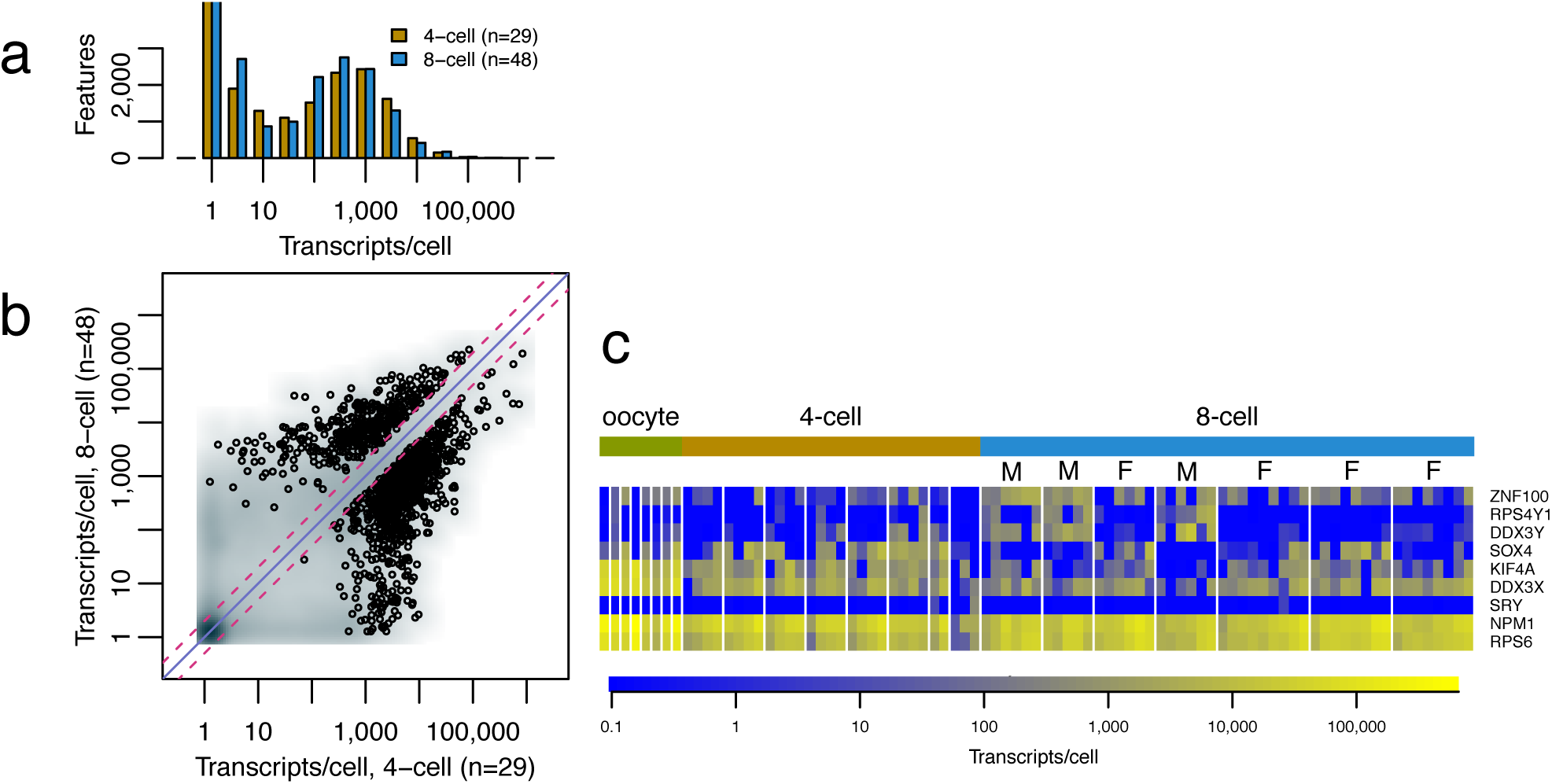
4- to 8-cell transition shows the major embryonic genome activation (EGA) a) The comparison of mRNA molecular count per cell showed reduction of transcripts in the 8-cell blastomeres compared to the 4-cell stage. b) Comparison of mRNA molecular count per cell per transcript on L185 library shows the dynamics in 4- to 8-cell transition with massive decrease of mRNA and major EGA at the same time. c) Expression patterns of sex-specific genes in the L185 library indicating paternal genome expression at this stage in development. Columns are cells separated by embryo. The individual cells within an embryo are sorted by transcript count/cell; with transcript-rich cells to the right. *ZFY* and *SRY* have been shown previously to be expressed before 2-cell stage by RT-PCR^37^. We detected additional sex-specific gene transcripts using STRT sequencing of single blastomeres, namely Y-specific *DDX3Y* and *RPS4Y*. When grouping cells based on the expression of *DDX3Y* and *RPS4Y*, the primate specific zinc finger gene *ZNF100* was also enriched in candidate male embryos, while *SOX4, KIF4A* and *DDX3X* were decreased. *NMP1* and *RPS6* were used as controls (Supplementary Data 2).

There has been much effort during the years to find polarity and cell fate commitment in 8-cell embryos. Changes in transcriptional profiles in the mouse have indicated transcription factors as the key players^22-24^. However, a recent study using single cell cDNA microarrays could not detect cell fate commitment at human 8-cell stage^3^.

When investigating polarity from the STRT sequencing data, we noticed positive skewness of transcripts/cell distribution within 8-cell embryos (Fig. 5a). In further analysis using two mouse primary cell populations^5^, we observed different distributions of skewness between the two mouse cell types and that of human 8-cell blastomeres, the blastomeres showing significantly larger skewness. Such a positive skewness value suggests the existence of a few transcript-rich blastomeres per embryo (P=0.01953 by Welch two sample t-test; Fig. 5b).

**Figure 5.**
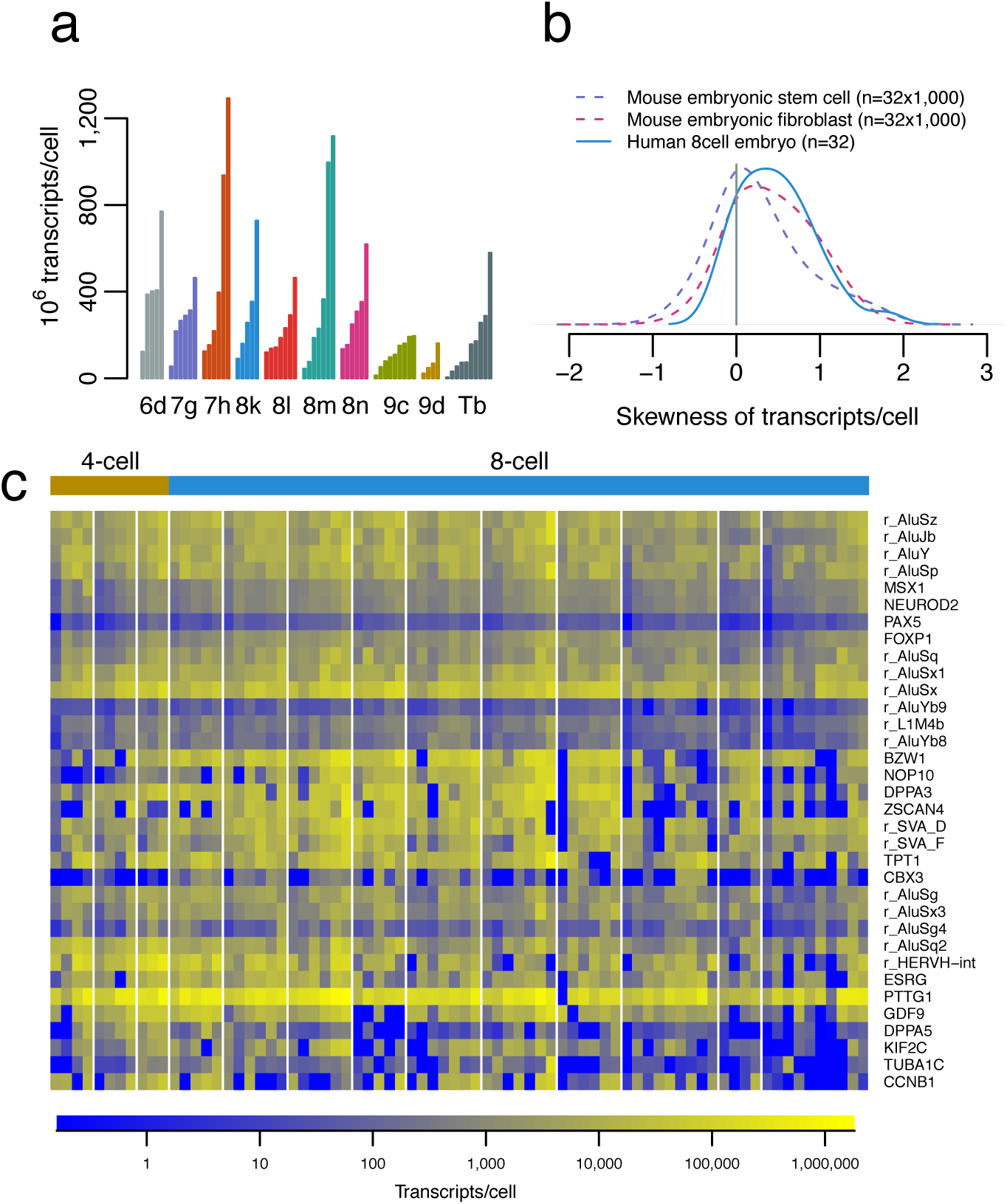
Differences in individual 8-cell blastomere transcriptomes. a) A variation in transcript count per individual blastomere was detected in 8-cell embryos on the L186 library. Individual blastomeres are colored by embryo in the figure. This shows that each embryo contains one or two transcript-rich blastomeres. ‘Tb’ stands for 10 cell embryo, the stage just previous to embryo compaction. b) Skewness distributions in human 8-cell stage embryos and equivalently sampled mouse embryonic stem cells and fibroblasts. The positive skewness property observed in 8-cell embryos was also seen in 4-cell embryos (Supplementary Fig 1). Our result supports the observation of different developmental potencies within individual blastomeres. Also, two independent studies report poor success rates in stem cell derivation and blastocyst formation when single 4-cell stage blastomeres were used^38,39^. c) A total of 155 genes and repeats were enriched in the transcript-rich blastomeres in all 8-cell embryos. Representative genes and repeats are shown in the heat map encompassing transitions from oocyte-to 8-cell stage blastomeres, on the L186 library. Columns are cells separated by embryo. The individual cells within an embryo are sorted by transcript count/cell; with transcript-rich cells to the right.

By comparing the two most transcript-rich blastomeres with the two most transcript-poor blastomeres in each 8-cell embryo a polarized pattern of 155 accumulated transcripts and repeats in the transcript-rich blastomeres was revealed (Fig. 5c and Supplementary Data 2). Interestingly, five *PRAMEF* genes (4, 5, 6, 9/15) recognized as DUX4 targets at 4-cell stage (Fig. 3c), were found enriched in the polarized 8-cell stage blastomeres. In particular, marker genes involved in pluripotency and stemness (*DPPA5, DPPA3* known as *STELLA, CBX3, TPT1, ZSCAN4, BZW1, PAX5, ESRG*), in telomere elongation (*NOP10*), transcription factors (*NEUROD2, FOXP1, PAX5*) and development (*GDF9, PAX5, KIF2C, NEUROD2, MSX1, FOXP, TUBA1C* and others) were accumulated in the transcript-rich 8-cell blastomeres. The *de novo* expression of Y-chromosomal genes was independent from transcript abundance between blastomeres. Within the group of genes accumulated in the transcript-rich blastomeres, 12 genes were involved in cell cycling, among them *Cyclin B1* (known as *CCNB1*), involved in the G2 to M transition^25^, and *Securin* (also known as *PTTG1*), involved in preventing premature chromosome separation^26,27^. The cell cycle control in the 8-cell blastomeres is mainly unknown, however, a recent report suggests that blastomeres overexpress cell cycle drivers and underexpress G1 and G2 checkpoints^28^.

Furthermore, among the accumulated features in the transcript-rich blastomeres we detected 17 repeat elements out of which 13 were Alus (Fig. 5c). The vast majority (9 out of 13) of these repeats were from the ancient AluS-type of repeats^29^. The Alu repeats (SINES, Short Interspersed Elements) are restricted to the primate lineage and are composed of a RNA polymerase III promoter and a polyA tail. Alu elements are shown to alter epigenetic regulation of gene expression^30^ and to contribute a large number of transcription factor binding sites^31^. Moreover, Alu repeats contain significant numbers of hormonal binding sites, such as Retinoic Acid Response Elements (RAREs)^32^. Possible regulation of these elements through retinoic acid receptors (RARs) is appealing since retinoic acid (RA) has been shown to affect many aspects of embryonic development in a wide variety of organisms^33^. In addition, PRAMEFs have been shown to interact with RARs and repress the transcription of RA target genes^34^. Since RA signaling is important for differentiation and early development, the expression of *PRAMEF* gene family members may imply an important human-specific early developmental function of these genes. We propose that the accumulation of *PRAMEF* and other markers in the transcript-rich 8-cell blastomeres indicate that these particular blastomeres are destined to become the inner cell mass (ICM).

Taken together, we addressed fundamental questions on human preimplantation development and uncovered transcriptome characteristics implicating DUX4 as an early embryonic activator expressed in the zygote. Moreover, we detected signs of embryo polarity already at the 8-cell stage. Certainly, mRNA expression does not accurately predict protein abundance or activity. However, the actual mRNA content at the earliest developmental stages needs to be understood at single cell resolution. Knowledge of the complexity in gene expression changes within single cells during the different embryonic stages is critical in understanding basic molecular mechanisms and functions in normal and subsequently abnormal preimplantation development. In summary, our analysis provides a vital foundation for future in-depth studies of early human development.

## Acknowledgements

We thank the anonymous donors of cells for this study. This study is the result of balanced contributions by the Hovatta, Kere and Linnarsson laboratories. We thank Manuel Pensis, Roberta Pecorari and Giammarco Momi from Procrea for zygote thawing procedures. This work was supported by the Karolinska Institutet Distinguished Professor Award to JK. The Swedish Research Council and ALF (Stockholm County and Karolinska Institutet) to OH and Åke Wiberg and Magnus Bergvall foundation to VT. The computations were performed on resources provided by SNIC through Uppsala Multidisciplinary Center for Advanced Computational Science (UPPMAX) under Project b2010037.

## Author contribution statement

V.T, S.K. and J.K. designed the experiment and wrote the manuscript. S.K. analyzed all sequencing data. V.T. participated in the sample collection and performed all expression validations. L.V. participated in planning the experiments and writing the manuscript. L.A., M.S. and O.H. were responsible for embryo thawing and blastomere biopsies in Sweden. G.F. organized the sample collection in Switzerland and M.J. was involved in the planning. A.J. and S.L. managed the sequencing facility and contributed to the result interpretation. O.H., S.L. and J.K. conceived the study. All authors contributed to the final manuscript.

## Material and methods

### Ethical statement

This study was reviewed and approved by the ethics review boards according to the applicable law in Sweden (The Regional Ethics Board in Stockholm) and in Switzerland. All cells were donated by couples who previously had had infertility treatment by *in vitro* fertilization and whose cells not needed for treatment had been cryopreserved. The donations of cells were by informed consent; the donated cells were destined for destruction after legal storage time had been reached. Zygotes were collected in Switzerland (authorization CE2161 of the Ticino ethical committee, Switzerland) and dissolved in reaction buffer before transfer to Sweden for sequencing. Oocytes and embryos were collected in Sweden.

### Human embryo culture and blastomere isolation

Non-fertilized MII oocytes were collected from the IVF clinic and the zona pellucidas (ZP) were removed using acid Tyrode’s solution (TS) (5mg/ml, Sigma-Aldrich). The dezoned cells were put into lysis buffer in individual wells on a 96-well plate and thereafter immediately frozen on dry ice. Zygotes were frozen at the pronuclear stage (2PN) 18-20 hours after fertilization. After thawing, the ZPs were removed using TS and treated as above.

Cleavage stage embryos were frozen at 4-cell stage on day 2 after fertilization. After thawing, embryos were allowed to develop until 6-10 cell stage in a sequential culture system (G1/CCM medium, Vitrolife) at 37°C and 5% CO_2_ and 5% O_2_. Individual blastomeres were obtained from each embryo by laser-assisted biopsy. Briefly, the embryos were held with a holding pipette and a laser beam created approximately 50μm big holes in the ZP. Individual blastomeres were aspirated and put individually into lysis buffer and frozen for downstream applications. All embryos used for blastomere biopsy were of equal size with little or no cytoplasmic fragmentation.

### Single cell mRNA sequencing using the STRT method

In this study we applied single-cell tagged reverse transcription (STRT), a highly multiplexed method for single-cell RNA-sequencing^5^. Each well of the 96-well plate contained individual cells with 5 μl of lysis buffer and a different template-switching helper oligo with a distinct six-base barcode and a universal primer sequence (for a detailed protocol see Islam et al^40^). Reverse transcription reagents were added to generate first-strand cDNA and eight synthetic spike-in RNAs were added to each well. Total concentration of the spike-in RNAs was 2,234.8 molecules/well^5^. After cDNA synthesis, the 96 reactions were pooled, purified, and amplified by single-primer PCR in a single tube in order to reduce cell-to-cell amplification variation and the number of PCR cycles. The amplified samples were sequenced on the Illumina platform. Sequenced STRT reads were filtered, trimmed, mapped to human UCSC genome hg19 and known repetitive elements, and counted by features, which consists of gene models and the repetitive elements. A detailed description of the pre-processing will be published elsewhere, and the software is freely available from the authors. Cells were excluded from further steps in case of shallow sequencing depth (requirement was at least 100k reads/well, including barcode sequence), no spike-in reads or problems in the cells observed by visual inspections (ex. lysed).Transcripts/cell was estimated by total barcode-containing reads and total barcode-containing spike-in RNA associated reads. We tested differential expression by SAMseq^41^ with a modification for spike-in-based normalization; the modification is that re-sampling of all annotated reads to be equal spike-in reads (see Supplementary Note, also more detailed description and source code for the statistical test will be published elsewhere).

### Single cell mRNA sequencing using the Tang method

Zygotes were frozen at the pronuclear stage (2PN) 18-20 hours after fertilization. After thawing, the ZPs were removed using TS followed by three washes in PBS-BSA (1mg/ml) droplets. Each single cell were put into a 0.5 ml PCR tube containing 4.45 μl of freshly prepared cell lysis buffer according to the protocol by Tang et al^6^. In total, 12 single zygote libraries were prepared, whereof four were sequenced on the SOLiD platform version 4.

Cleavage stage embryos were frozen at 4-cell stage on day 2 after fertilization. After thawing, the 4-cell embryos were dezoned using TS followed by three washes in PBS-BSA (1mg/ml) and put into lysis buffer for downstream application. In total, four 4-cell embryos were collected for sequencing.

### Visualization of single cell RNA sequencing results on human genome

For visualization purpose, raw RNA sequencing reads were sorted by barcode (only for STRT), trimmed, and mapped to reference sequences, which consists of human genome UCSC hg19, human ribosomal repeating unit (GenBank: U13369) and the spike-in mRNA sequences, using TopHat^42^. For the visual inspections of transcription start sites using STRT reads, we used 5’-end position of the alignments. Read count was divided by the number of aligned sites, for example, one multiply aligned read at 10 sites were accumulated as 1/10 read at the 10 sites. We used UCSC Genome Browser^43^ for the genomic visualization, with the UCSC hg19 assembly.

### Transcript validation

In order to further validate the presence of full-length *DUX4* transcripts in zygotes, we used nested PCR on a single cell cDNA library constructed according to Tang 2010^6^. The libraries were amplified with the primer pairs, 14A∼174 followed by 16A∼175 covering the full length *DUX4* transcript^9^. After gel purification the fragments were cloned into pTOPO vector (Invitrogen) and sequenced with T3 and T7 primers by Sanger method.

